# Effects of fine-scale population structure on the distribution of heterozygosity in a long-term study of *Antirrhinum majus*

**DOI:** 10.1101/2020.08.20.259036

**Authors:** Parvathy Surendranadh, Louise Arathoon, Carina A. Baskett, David L. Field, Melinda Pickup, Nicholas H. Barton

## Abstract

Many studies have quantified the distribution of heterozygosity and relatedness in natural populations, but surprisingly few have examined the demographic processes driving these patterns. In this study we take a novel approach by studying how population structure affects both pairwise identity and the distribution of heterozygosity in a natural population of a self-incompatible plant *Antirrhinum majus*. We look at a measure of the variance in heterozygosity within a population, identity disequilibrium (*g*_2_), together with F_ST_ using a panel of 91 SNPs in 22,353 individuals collected over 11 years. We find that pairwise relatedness (F_ST_) declines rapidly over short spatial scales, and the excess variance in heterozygosity between individuals (*g*_2_) reflects significant variation in inbreeding. Additionally, we detect an excess of individuals with around half the average heterozygosity, indicating that some are due to selfing or matings between close relatives. We use two types of simulation to ask whether variation in heterozygosity is consistent with fine-scale spatial population structure. First, by simulating offspring using parents drawn from a range of spatial scales, we show that the known pollen dispersal kernel explains *g*_2_. Second, we simulate a 1000-generation pedigree using the known dispersal and spatial distribution and find that the resulting *g*_2_ is consistent with that observed from the field data. In contrast, a simulated population with uniform density underestimates *g*_2_, indicating that heterogeneous density promotes identity disequilibrium. Our study shows that heterogeneous density and leptokurtic dispersal can together explain the distribution of heterozygosity. Furthermore, our study highlights the limitations of making theoretical predictions from simulations that only assume simple density and dispersal distributions.

## Introduction

Spatial genetic structure is common in natural populations. For many organisms, gene dispersal and therefore relatedness are spatially structured, such that individuals closer in space are more likely to mate, and be more closely related, than individuals further apart [1], [2]. Such spatial population structure should reduce population mean heterozygosity relative to a well-mixed population, generating decreasing genetic similarity over geographic distance – known as isolation-by-distance [3]. Despite the ubiquity of these patterns in nature, the importance of demography and gene dispersal in determining both the spatial pattern and level of genetic variation has not been thoroughly explored. Furthermore, commonly used models of spatial structure – namely the island model, stepping stone model and models of continuous population structure – either assume demes or a uniform population density. However, natural populations are typically patchy, with heterogeneity in both the distribution and density of individuals. Patchy and heterogeneous spatial distributions within natural populations should result in spatial variation in inbreeding and, consequently, excess variance in heterozygosity. Despite this prediction, spatial variation in heterozygosity has rarely been examined in the population structure literature. Moreover, it is the interplay of heterogeneous density and dispersal that likely shapes the spatial structuring of genetic relatedness between individuals. This highlights the importance of understanding the factors (e.g., life history, demography, population structure) that contribute to shaping the full distribution of heterozygosity and relatedness in a spatially structured population.

Understanding the drivers of variation in inbreeding within populations is fundamental given its importance to genetic diversity and fitness. Quantifying variation in inbreeding and combining this with measures of fitness (or fitness proxies) makes it possible, in principle, to estimate inbreeding depression either through pedigrees [4], [5] or Heterozygosity-Fitness Correlations (HFCs). For HFCs, inbreeding depression is estimated by comparing proxy measures of fitness against heterozygosity, with the expectation that offspring from related individuals will have lower heterozygosity. Variance in inbreeding is therefore essential for HFCs to be detected [6]. In addition, variance in inbreeding is interesting *per se* because it depends on both demographic history (e.g., [7]) and mating system (selfing, partial selfing or outcrossing) [8]. For mating system, selfers vs. outcrossers can show contrasting levels of inbreeding, with different consequences for genetic diversity, depending on population history. Outcrossing species, with generally low levels of inbreeding, provide an opportunity to examine factors other than mating system variation that may affect inbreeding variation, and thus, variance in heterozygosity.

If there is variation in inbreeding between individuals, heterozygosity at different loci will be correlated. The covariance between loci in heterozygous state is termed identity disequilibrium (ID), by analogy with linkage disequilibrium, the covariance in allelic state between loci. ID can be calculated across individuals and divided by the square of the mean of heterozygosity to calculate the population statistic *g*_2_, which is a measure of variance in identity by descent [6]. For an outcrossing organism with fine-scale population structure, spatial patterns of density and mating could have strong effects on the degree of mating with related individuals, and thus affect identity disequilibrium and *g*_2_. Furthermore, as sessile organisms, mating and offspring dispersal in plants are mediated by external vectors (pollinators and seed dispersal mechanisms) [9]. Consequently, the shape of the distribution of dispersal of both pollen and seed will also have an impact on *g*_2_. If the sources of variation in inbreeding are better understood, we may be able to combine *g*_2_ with other statistics of population structure to improve inferences about demographic history [10], [11]. Thus, *g*_2_ can be utilized to learn about the effects of population structure on the distribution of heterozygosity and pairwise identity.

For over a decade, we have sampled a population of the self-incompatible plant *Antirrhinum majus*, the long-term aim being to build a pedigree which will allow us to estimate fitness and dispersal directly. Through that project, we have collected an exceptionally large sample of marker genotypes, which enables a powerful test of whether the observed density and dispersal in this population can account for both the decay of pairwise relatedness with distance, and for the distribution of heterozygosity across individuals. First, we verify that there is excess variance in heterozygosity, which reflects an underlying variance in inbreeding. Second, to understand the role of spatial patterns of dispersal in generating variance in heterozygosity, we compare the empirical distribution of heterozygosity with that of offspring from simulated matings where parents were drawn from different dispersal scales. Third, we ask whether heterogeneous population density promotes variation in inbreeding, by comparing simulated pedigrees conditioned on uniform density versus on the observed locations of plants. Taken together, addressing these questions provides insight into the underlying drivers of spatial structure and provides novel ways to study the effects of mating patterns and demography in nature.

Throughout this paper, it will be important to distinguish between identity by descent (IBD) and identity in state. We denote the probability that two genes are identical by descent by *F*; this is defined relative to an ancestral reference population, and can in principle be calculated from the pedigree that descends from that population, independent of the actual allelic state. What we observe are SNP genotypes; the two homologous genes in a diploid individual will be identical in state if the genes are identical by descent, or if the ancestral genes carried the same allele. Thus, probabilities of identity by descent (*F*) can be estimated from observed identities in state. We denote the heterozygosity at locus i in a particular individual by h_i_, with h_i_=0 if the genes are identical in state, and h_i_=1 otherwise. The mean heterozygosity of an individual is the average of h_i_ over n loci, denoted 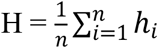.

## Methods

### Study system

*Antirrhinum majus* is a self-incompatible, hermaphroditic, short-lived perennial herb native to the Iberian Peninsula. It has a seed bank with most individuals’ parents recorded 3-4 years before they are sampled (D. Field, unpublished data). It grows in a variety of microhabitats with relatively bare soil or frequent disturbance, including rail embankments, rocky cliffs, and regularly mowed roadsides. Our study includes two “subspecies” that differ only in flower color: *A. majus pseudomajus* has magenta flowers and occurs in northern Spain and south-western France, including the Pyrenees. *A. majus striatum* has yellow flowers and a smaller range, encircled by *A. m. pseudomajus*. The subspecies are parapatric; narrow clines with intermediate color hybrids form wherever they meet, and there is no evidence for post-zygotic reproductive barriers [12]. We focus on such a hybrid zone in the Vall de Ribès, Spain [13], where we have collected demographic data annually since 2009. Across nearly all of the genome, there is little divergence within our study area between plants with different flower color, except for limited regions associated with floral pigmentation, which show steep clines [14]. Thus, the study area can be considered as a single population for studying neutral genetic variation.

### Field sampling

Genetic samples were obtained annually from 2009-2019 from every accessible flowering individual in ~5 km stretches of two parallel roads that cross the Vall de Ribès, dubbed the “lower road” (GIV-4016; ~1150 m elevation) and “upper road” (N-260; ~1350 m) (Fig. 1). We also sampled along small side roads, railroad embankments, rivers, and hiking trails; density was very low away from these disturbed areas. In some years, we were limited to genotyping only in the core area, ~1 km along each road. The total genotyped sample summed over the eleven years is 22,353 plants, ranging from ~750 plants in the smallest year (2018), to ~5500 plants in the largest year (2014). Eighteen percent of individuals were sampled in more than one year. Sampling was conducted during peak flowering (early June to mid-July). Each year there were fewer than 100 visible but inaccessible plants; consequently, we estimate that we found the majority of individuals in the sampled area.

**Figure 1:**
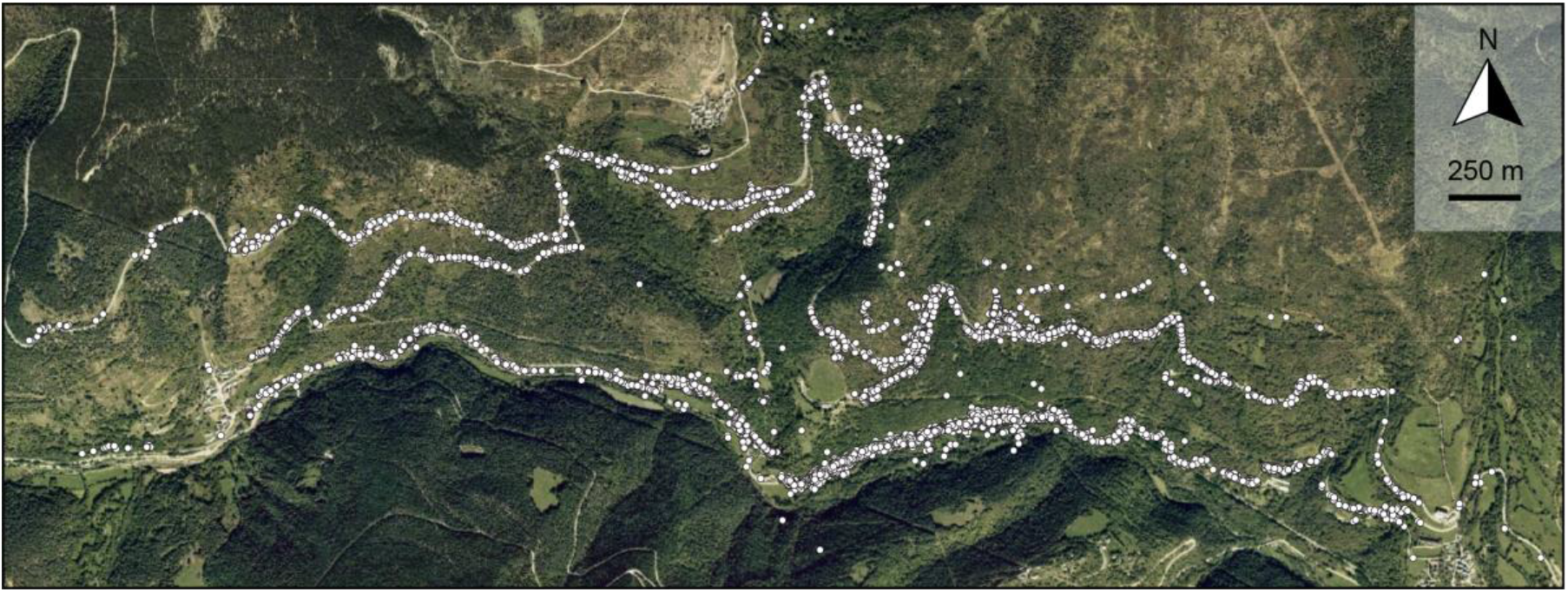
Distribution of *A. majus* in Vall de Ribès, Spain from the years 2009 to 2019, each individual is shown as a white circle.

For each plant, we collected leaf samples for genotyping, and recorded spatial locations with GeoXT handheld GPS units (Trimble, Sunnydale, CA, USA). These devices are accurate to within 3.7 m, determined by comparing samples that had been inadvertently recorded twice in the field (individuals with similar geographic location and near-identical genotypes, allowing for SNP errors). Leaf samples were refrigerated upon return to the field station, and dried in silica gel within three days.

### SNP panel

Previously, a panel of 248 SNPs spread throughout the genome was designed for the focal population (see methods in [15]). We follow these methods but include an additional five years of data (2015-2019) and use a subset of 91 SNPs; the mean sample size per SNP was 21,212, or ~95% of the total. (see Supplemental Material 1.1 (SM1.1) for SNP filtering methods). We imputed the ~5% missing genotypes for each SNP by randomly assigning genotypes according to the population-wide allele frequencies at each marker.

### Isolation by distance

The panel of 91 SNPs was used to calculate F_ST_ and isolation by distance, both of which relate to the mean heterozygosity. F_ST_ is defined as the average identity by descent among individuals within subpopulation, F_S_, relative to the total population, 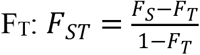 [16].

These identities are estimated from SNP genotypes since we do not have the full pedigree. Two genes will have a different allelic state only if they are not identical by descent, and if they derive from different alleles in the ancestral population. Given overall allele frequencies p+q=1, the expected heterozygosity 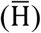 of offspring from parents whose genes have a probability of identity by descent *F* is 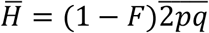, where 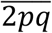 is an average over loci. Thus, there is a direct relation between F_ST_ and the mean heterozygosity: 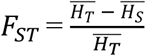. We use this relation to compute F_ST_ for this dataset. (Note that here, H is the probability of non-identity in state, which depends on the SNP genotype. The subscripts S and T refer to the specified quantity within subpopulation and total population, respectively). A subpopulation is defined as individuals within 20m.

Isolation-by-distance – the decay of genetic similarity with geographic distance – can be observed by measuring pairwise relatedness between individuals. If individuals are separated by a distance r, then pairwise relatedness can be calculated as an extension of F_ST_ (which we refer to as pairwise F_ST_) by setting F_S_ to be the probability of identity by descent and, correspondingly, 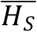 to be the probability of non-identity in state between genes which are at a distance r apart. This formulation is used to estimate F_ST_ between every pair of individuals relative to the total population, as a function of their geographic separation. Pairs of individuals are binned into distance classes of 20m each and the average F_ST_ and the distance corresponding to each bin is calculated. This was done for every year from 2009 to 2019, and the average calculated.

### Variation in inbreeding

To describe the variation in heterozygosity in the *A. majus* population, we calculated multilocus heterozygosity, denoted here by H, pooling across all years. This is defined as the fraction of heterozygous loci in an individual: 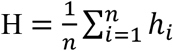, where *h_i_* = 0 or 1, according to whether locus *i* is heterozygous, and there are *n* genotyped loci. In this system “generations” cannot be clearly defined because of seed dormancy and perenniality. However, pooling data across years only reduced H by 0.08%.

We observed an excess of individuals with around half the mean heterozygosity (see Results). To check whether the pattern was consistent with rare selfing, we compared the likelihood of a single Gaussian to a mixture of two Gaussian distributions, one with the observed mean and variance and the other with half its mean and variance.

The variance in individual heterozygosity consists of two components. The first is due to the variance in whether an individual locus is heterozygous, and decreases in proportion to the number of SNP, *n*: it equals 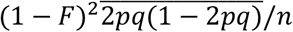. The second is due to covariance in heterozygosity between loci, which is termed the identity disequilibrium (ID). For a given pedigree, unlinked genes flow independently. Thus, heterozygosity is independent across unlinked loci, and so this second component is proportional to the variance in inbreeding across individuals, var (*F*). The first component can be estimated from the allele frequencies, or simply by shuffling the data across loci, to eliminate ID. The excess variance is then proportional to the variance in F across individuals, and is measured by the statistic *g*_2_:

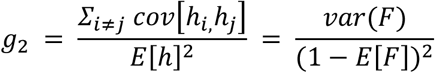

(from Eq. 1 in [6]). Here, cov[*h_i_*, *h_j_*] is the ID between loci i and j, and the sum over all distinct i,j is the excess variance in H due to ID. Dividing by the square of the mean heterozygosity E[h]^2^ eliminates dependence on allele frequency, such that *g*_2_ estimates the variance in *F* across individuals.

To describe the variance of inbreeding across individuals, we first check if the variance in the distribution of individual heterozygosity is significantly greater than the average variance obtained from 100 replicates. This was done by reassigning heterozygous status randomly across loci within individuals, which would eliminate correlations between loci generated by ID. We then computed *g_2_* using the g2_snps function from the R package InbreedR (in R version 3.6.1 [17]), which implements a modified formula for large data sets to estimate *g_2_*, and provides confidence intervals via bootstrapping to account for the finite number of individuals sampled [18]. We decomposed ID into components due to linked and unlinked SNPs by comparing correlations of H for all individuals to those with low H, at several scales: across all pairs, within linkage groups, and between adjacent SNPs (SM1.2: Table S1).

### Effects of pollen dispersal on heterozygosity

With isolation by distance, the distribution of heterozygosity is expected to depend on the distance between parents: heterozygosity of offspring from nearby parents will have a lower mean and higher variance compared to offspring from distant parents. To test this prediction, we simulated offspring using all field individuals as mothers and choosing fathers from a given distance away (detail in SM1.3). Then, we compared the distribution of H between the field data and offspring simulated from matings with three models of pollen dispersal: the nearest neighbor to the mother, a Gaussian distribution (σ = 300 m), and a leptokurtic dispersal kernel sampled from 1463 empirical measurements of pollen dispersal, estimated as the distances between assigned parents (electronic supplementary material; D. Field, unpublished data). The genotype of the offspring was assigned using Mendelian inheritance, either without linkage between markers, or using the known linkage map (electronic supplementary material; courtesy of Yongbiao Xue, Beijing Institute of Genomics). Including linkage did not substantially change results, so we mainly show results for simulations without linkage. We compared distributions, means, and variance of H using Kolmogorov-Smirnov tests, t-tests, and F-tests, respectively. For the leptokurtic pollen dispersal simulation, we checked for an excess of low-heterozygosity individuals generated by mating between close relatives by asking whether a mixture of two Gaussian distributions is more likely than a single Gaussian distribution.

### Heterozygosity in a simulated spatial pedigree

In order to compare the actual distribution of heterozygosity with that expected for a spatially structured population, we simulated a continuous two-dimensional population, conditioned on the known locations of the individuals and the empirically measured seed and pollen dispersal distances (electronic supplementary material; D. Field, unpublished data), using Mathematica 12.0 [19]. Our simulation differs from commonly used models (e.g., island [16], stepping stone [20] and continuous Wright-Malécot model [3], [21]) in that we include heterogeneity in density by specifying actual locations to determine relationships in the pedigree. We only provide a proof-of-principle, by asking whether a plausible model of spatial structure can explain the observed heterozygosity. We do not include all features of the actual population – in particular, we extrapolate by repeatedly sampling ten years of spatial distributions; we ignore linkage; we simplify the self-incompatibility system; and we assume an annual life cycle (no perenniality or seed bank). Thus, our simulation parameters should be seen as “effective” values, analogous to the traditional Ne. Additionally, we also validated our simulation against analytical results for a panmictic population and stepping stone model (SM1.4: Fig. S5).

First, however, we simulated a population with uniform density (the continuous Wright-Malécot model) as a null model, to compare expected heterozygosity with and without heterogeneous spatial structure. We simulate a region of ~1.1 x 1.8 km that was sampled consistently in the *A. majus* focal population (SM1.4: Fig. S3). Locations were assigned by randomly sampling N points from a uniform distribution each generation, for 1000 generations. Genetic diversity is shaped over the coalescent timescale (2N_e_, ~170,000 generations in *A. majus* [14]), which is far longer than the 1000 generations that we simulate. However, we are concerned here with the *local* population structure that determines the variation in inbreeding amongst individuals within an area of a few km^2^, which will equilibrate rapidly. The spatial pedigree was generated by choosing parents for each individual according to a backwards dispersal distribution measured empirically. The seed and pollen dispersal distances are estimated respectively as the distance between offspring and nearest parent (assumed to be the mother) and between parents (electronic supplementary material; D. Field, unpublished data). For every offspring, the mother and father are chosen from randomly drawn distances from the seed and pollen dispersal distributions. To choose a parent from a distance r, 6 points are assigned randomly on a circle of radius r centred at the focal individual and the nearest individual to each of them are found. The closest individual to any of these points is then chosen as the parent. The accuracy of our algorithm is verified by comparing the specified and realised seed and pollen dispersal distributions for the simulated pedigrees (SM1.4: Fig. S6, Table S4). The same procedure is repeated for the father, taking the mother as the starting point. Since *A. majus* is self-incompatible, the mother and father are not allowed to be the same individual.

Once the spatial pedigree is generated, 10 replicate sets of genotypes are assigned by dropping genes down the pedigree, starting with equal expected frequencies of both alleles at each of 91 loci. In fact, one could start with any initial frequencies, since F_ST_-like measures are independent of them. Population size was adjusted so that F_ST_ matched the empirical data for the simulated sampling area.

Next, we simulated a population with realistic heterogeneous spatial structure by using the individual locations available for the years 2009 to 2019 in the *A. majus* focal population (SM1.4: Fig. S4). There were fewer individuals from 2017-2018, so these were merged, giving distribution data for 10 time points. We randomly sample from the ten consecutive time points, and repeat for 100 cycles, thus iterating for 1000 generations. We sub-sample from these locations to maintain a constant population size (N). If N is greater than the number of plants available in a given time point, say k, all k plants are first included and the remaining N-k locations were re-sampled from the same time point, displaced at a random angle on a circle of radius 3m to avoid having plants in the same location. This naïve approach allows us to simulate a spatial structure that is realistic over at least small scales. We then generated a pedigree following the procedure used for the uniform population, again adjusting population size to match the empirically observed F_ST_. Ten replicate sets of genotypes were run for each of five replicate pedigrees.

Patterns of isolation by distance, heterozygote deficit (F_IS_) and identity disequilibrium were compared between the two simulation types and the field data (calculated from the simulated sub-area of the field site). As the fitted population sizes were large (see Results), obtaining direct estimates of identity by descent and thus F_ST_ from the pedigrees was not feasible. Instead, F_ST_ was obtained for a pedigree as the average of replicate genotype sets generated from that pedigree. F_IS_ was calculated from the observed and expected heterozygosity. Values of *g_2_* were calculated for each replicate from each pedigree using InbreedR (in R version 3.6.1 [17]).

## Results

### Isolation by distance

The average pairwise F_ST_ was calculated each year for individuals separated by different distance classes and then averaged across years. Pairwise relatedness (F_ST_) between individuals decreased rapidly with geographic distance, showing isolation by distance (Fig. 2A). If we consider individuals within geographic separation of 20m, the average F_ST_ over the eleven years is 0.0244; however, this figure is an average over a quantity that depends strongly on distance. The sharp decline in pairwise identity over short spatial scales corresponds precisely to a rapid increase in H with distance between parents (SM1.3: Fig. S1), since heterozygosity is determined by the probability of identity by descent between the genes from each parent. Note that over large separations (>1Km), pairwise F_ST_ values are necessarily negative, because distant individuals are less closely related than the average for the whole population.

**Figure 2.**
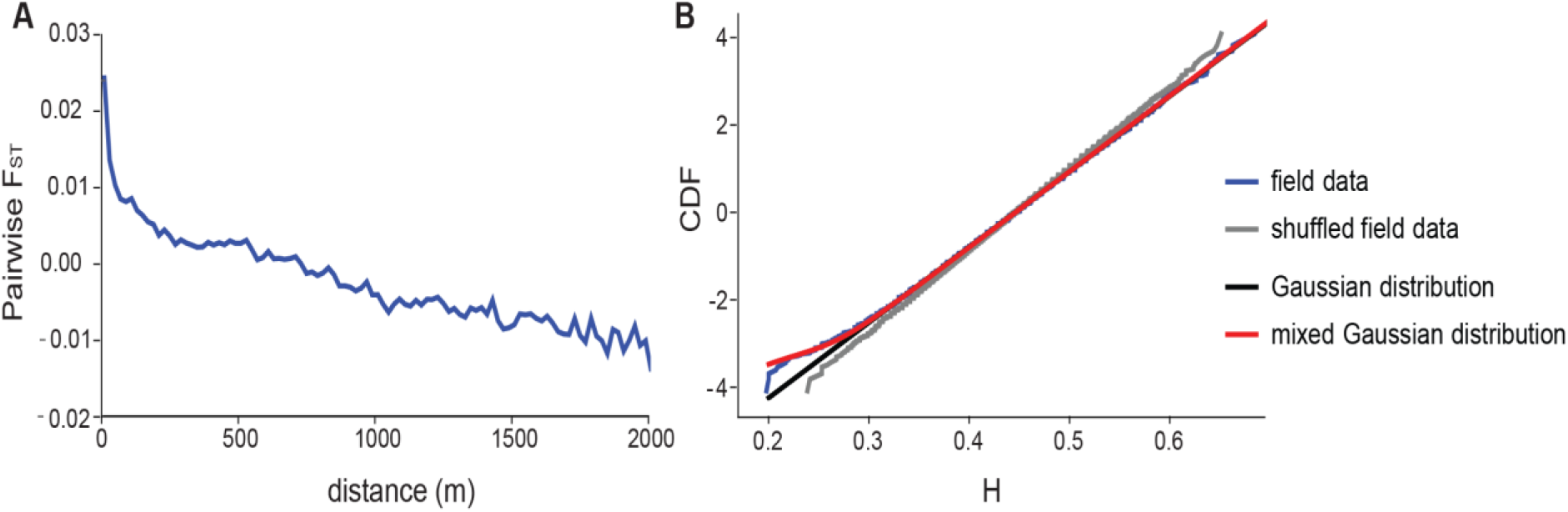
**A**: Pairwise relatedness (F_ST_) between individuals decreases rapidly with geographic distance showing isolation-by-distance in the field data. **B**: Probit transform of the cumulative distribution function (CDF) of the distribution of individual heterozygosity (H). A Gaussian appears as a straight line on a probit scale, and the y-axis is the number of standard deviations of the standard normal distribution.

### Variation in inbreeding

Excess variance in the distribution of individual heterozygosity (H) in the field data shows that there is variance in inbreeding in the population (Fig. 2B). Furthermore, there is an excess of individuals with around half the mean heterozygosity (i.e., with H~0.22, rather than 0.44; Fig. 2B, blue, lower left). These might be due to a low rate of selfing. Indeed, a mixture between two Gaussian with means ~ 0.22 and 0.44, and variances in the same ratio, fits significantly better than a single Gaussian (Fig. 2B, compare red and black to blue) with an increased likelihood of 11.3. However, we shall see in the next section that this excess is also consistent with matings between close relatives, without the need to invoke a breakdown in self-incompatibility.

To examine whether the observed distribution of heterozygosity is significantly different to a null distribution, we compared the field data with heterozygous values shuffled across individuals, which eliminates identity disequilibrium (ID) by removing correlations between loci. We found greater variance in heterozygosity in the observed compared to the randomly shuffled field data (Fig. 2B, gray). For both data sets, the mean heterozygosity (0.44602) necessarily remains the same, but the observed variance in the field data (var(H) = 0.00336) was significantly higher than the average variance in 100 shuffled replicates (mean var(H) = 0.00282, s.d. 0.000029). This excess variance between the observed and shuffled data implies that the mean standardized ID is *g_2_* = 0.0029 (95% CI: 0.0026-0.0033), representing a significant variance in inbreeding between individuals.

The overall ID, as measured by *g*_2_, is due to correlations in heterozygosity between all pairs of loci, most of which are unlinked. We expect stronger correlations between linked loci, because relatives will share blocks of genome. We found that the mean covariance in heterozygosity between SNP on the same linkage group is substantially stronger than the overall mean (0.00265 vs. 0.00056). If we restrict attention to those individuals with H<0.3, we find that the covariance in heterozygosity between SNP on the same linkage group is still higher (0.00649), as expected if close relatives share long blocks of genome IBD (SM1.2: Table S1).

### Effects of pollen dispersal on heterozygosity

The heterozygosity of simulated offspring depends on distance between their parents, with a rapid increase in mean H with distance (SM1.3: Fig. S1). We compared the observed distribution of heterozygosity with three alternative scenarios for pollen dispersal. There was no significant difference between the mean and variance of heterozygosity between the field data and offspring simulated from the observed leptokurtic dispersal (electronic supplementary material). However, the mean and variance of heterozygosity differed between the field data and simulated matings with either nearest neighbors, or with Gaussian dispersal (Fig. 3A, SM1.3: Tables S2-S3). On the other hand, compared to the field data, all three dispersal schemes differed in the distribution tail as assessed by Kolmogorov-Smirnov tests (SM1.3: Table S3). These comparisons were made for a single replicate, but because each involves 22,353 individuals, there was little variation in the mean and variance between replicates.

**Figure 3.**
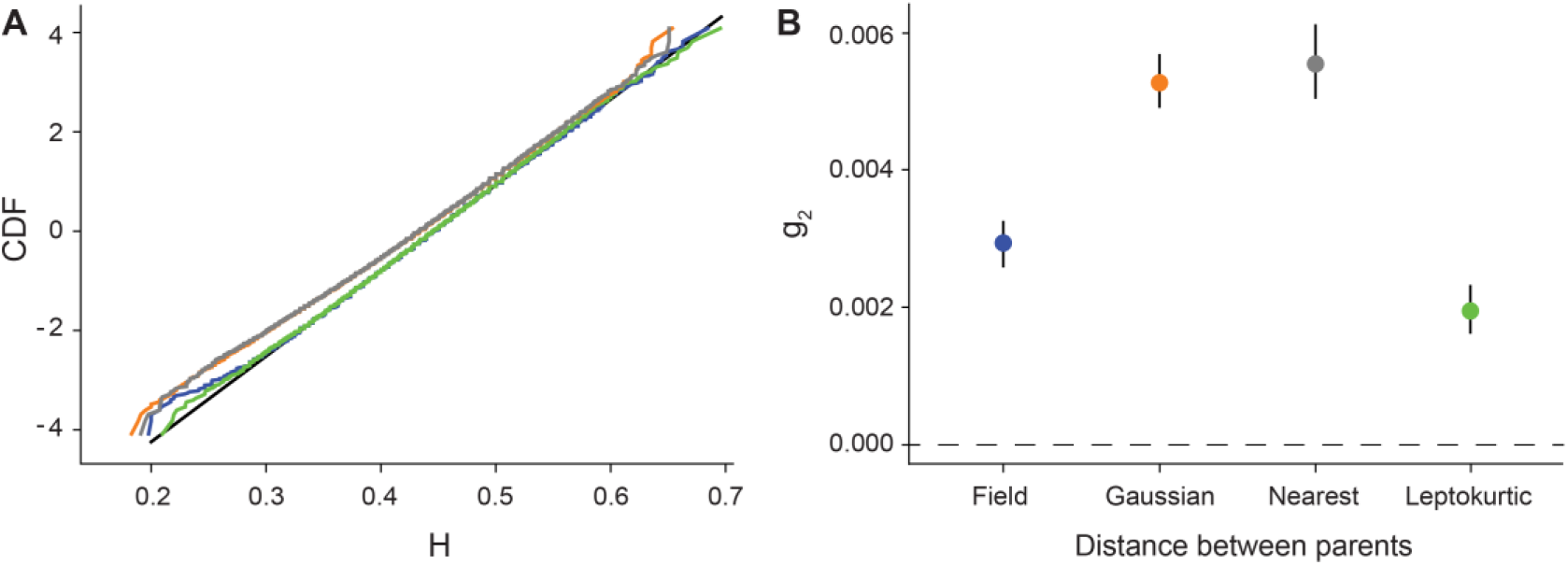
**A**: Probit transform of the CDF of multilocus heterozygosity, H, for the field data (blue) versus a single replicate of offspring simulated from Gaussian pollen dispersal (orange), nearest neighbor matings (gray), and leptokurtic pollen dispersal (green). A normal distribution (black) with the same mean and standard deviation as the field data is included for comparison **B**: Identity disequilibrium for the same data as above indicating mean and 95% CI.

We next examined deviations in the left tail of the distribution, where an excess of low heterozygosity individuals might arise from selfing or from matings between close relatives. We focused on the leptokurtic dispersal curve, which was the distribution closest to the field data. We estimated the increase in likelihood between fitting a single versus mixed Gaussian distribution (see “Variation in inbreeding”) for 100 replicate simulations. We found that the mixed Gaussian was a better fit than a single Gaussian, with an increase in log likelihood greater than 2 for 69 of 100 replicates. The estimated fraction of putatively “selfed” individuals was 0.00043, averaged over replicates, which is about half the estimate from the actual data, 0.00086. In comparison, only 4/100 replicates gave higher estimates than that observed (SM1.3: Fig. S2). This suggests that the excess of individuals with low heterozygosity can to a large extent be explained by matings between relatives under leptokurtic pollen dispersal. Nevertheless, there is a marginally significant excess of such individuals, with twice as many being seen as expected from our simulations. There is considerable variation in fit between replicates, simply because deviations in the tail involve few individuals

The coefficient *g_2_* reflects excess variation due to identity disequilibrium, and showed similar patterns as the variance in H. Here, we found no significant difference between *g_2_* from field data and offspring from simulated matings with leptokurtic pollen dispersal. However, *g_2_* from Gaussian and neighbor matings were 80% higher than *g_2_* from field data and leptokurtic matings. This nominally represents a significant difference given that the 95% confidence intervals between these groups do not overlap (Fig. 3B). However, as we discuss below, these confidence intervals only include sampling error, and not the additional variance due to random evolutionary realizations.

### Heterozygosity in a simulated spatial pedigree

In the previous section, we simulated offspring across one generation. To examine if the observed heterozygosity is consistent with a spatially structured model, we simulated a pedigree over 1000 generations, conditioned on the locations of individuals observed over ten years, repeated over 100 cycles. The realized seed and pollen dispersal matched the empirical seed and pollen dispersal distribution for both density types (SM1.4: Fig. S6, Table S4). We required N = 15500 individuals for the heterogeneous density model and 40000 individuals for the uniform density, in order to match the observed F_ST_ ~ 0.022 calculated over a 20m scale from the simulated sub-area of the field site (SM1.4: Table S5). Up to distances of 1km, the decline in pairwise identity with distance matched between the field data and the five replicate pedigrees simulated with heterogeneous density (Fig. 4A, SM1.4: Fig. S7A). High variation among replicates suggests that many more SNPs would be needed to match the pattern from the pedigree (SM1.4: Fig. S7B); moreover, linkage would increase this variance to some extent. We also compared the pattern of isolation by distance from the field data to that from the pedigrees generated for both the heterogeneous and uniform density scenarios (Fig 4B, SM1.4: Fig. S8); the heterogeneous density is a much better fit than the uniform density (SM1.4: Table S5).

**Figure 4.**
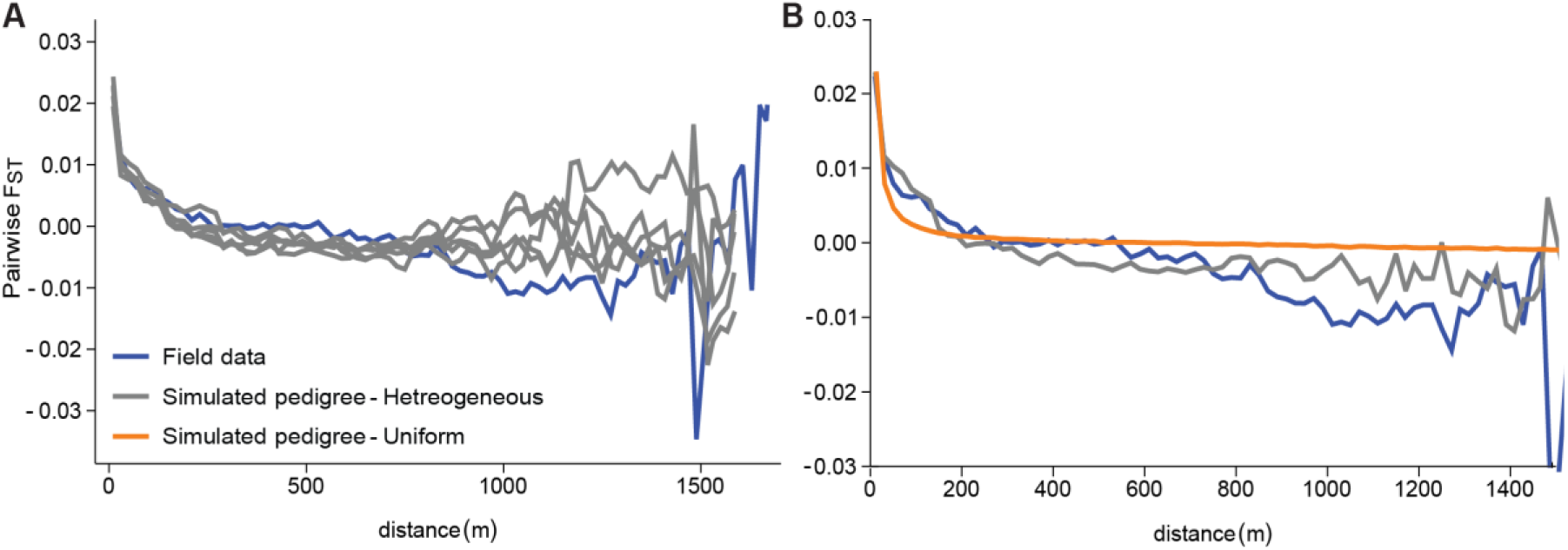
**A**:. Isolation by distance compared between the field data (blue) and five replicate simulated pedigrees (gray) based on a heterogeneous population density. **B**: Isolation by distance from the field data (blue) compared between the simulated pedigree with a heterogeneous (gray) and uniform (orange) population density.

Identity disequilibrium (*g*_2_) estimates from the genotypes from pedigrees simulated with heterogeneous density showed substantial variation between the five simulated pedigrees, and between the ten draws of 91 SNPs from each pedigree (Fig. 5). The average *g*_2_ estimated from the five pedigrees (each with 10 replicates) is 0.00264, which is consistent with the observed mean annual *g*_2_ from the field of 0.00262. On the other hand, when assuming a uniform density, the average *g*_2_ of 0.00171 is significantly lower than the field data. Note that the confidence limits for the field data, generated by InbreedR, only include error due to sampling a limited number of individuals. These errors do not account for sampling a limited number of SNPs, or the random variation between evolutionary realizations (see Discussion).

**Figure 5.**
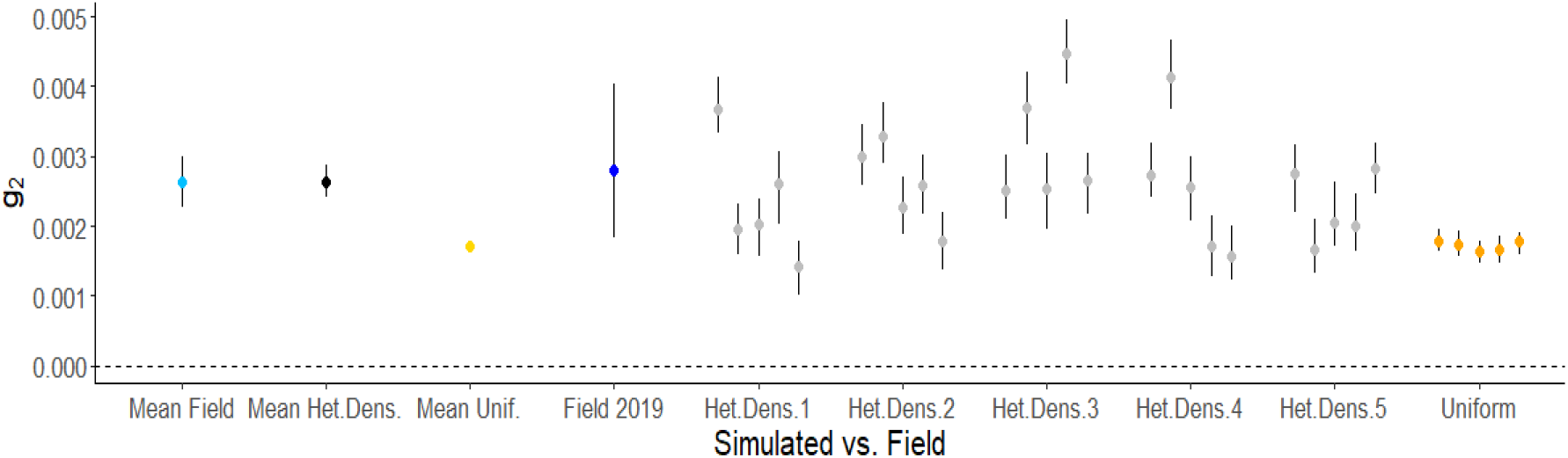
Identity disequilibrium (*g*_2_) calculated from field data versus simulated pedigrees. Five of ten replicates per pedigree are shown (gray: heterogeneous density, with five simulated pedigrees; orange: uniform density, with one simulated pedigree). Mean from the field (light blue) is across 2009-2019, while mean from the heterogeneous (black) and uniform (yellow) simulations is across all replicates. The final year of field data (dark blue) is comparable to *g*_2_ calculated from the final year of pedigree replicates (gray and orange).

## Discussion

An enduring problem in evolutionary biology is understanding how demographic processes, such as heterogeneous density and dispersal, interact with spatial structure to determine the distribution of heterozygosity within populations. In this study of a long-term dataset, including more than 20,000 plants sampled over 11 years, we combine field data and simulations to address questions central to understanding how demography can influence patterns of heterozygosity. Namely, can we predict the distribution of heterozygosity for an outcrossing species from key demographic parameters? To address this question, we first confirmed that there was significant correlation in heterozygosity between markers (*g_2_*, a measure of identity disequilibrium), which implies variation in inbreeding. By simulating offspring from matings between geo-referenced, genotyped individuals, we show that the mean heterozygosity increases, and the variance of heterozygosity decreases, with increasing distance between parents; strikingly, these changes occur over very short scales (~10m, SM1.3: Fig. S1). We found that the observed distribution of heterozygosity is consistent with the known leptokurtic distribution of pollen dispersal. We also simulate the population over 1000 generations using the actual seed and pollen dispersal kernels, and the observed heterogeneous density. We found that this model matches the observed identity disequilibrium, whereas a null model with uniform density substantially underestimates the observed patterns. Moreover, our results also highlight the limitations of making theoretical predictions from simulations that only assume simple demographies. Taken together, our findings highlight the potential for using the observed demography to explain the distribution of diversity, and specifically the variance in inbreeding in spatially continuous populations.

Variation in heterozygosity within populations provides the potential for selection to reduce the frequency of less fit, inbred individuals. The association between inbreeding and fitness is often tested through Heterozygosity-Fitness Correlations (HFC), which quantify inbreeding depression in natural populations by correlating measures of fitness with heterozygosity [6]. Many studies that test for HFCs find that the excess variation in heterozygosity, *g_2_*, which arises from identity disequilibrium, is low and rarely significant [22]. In our study, we estimate a significant *g_2_* of 0.0029 (95% CI: 0.0026-0.0033). Although low, this estimate is of the same order as most of the *g_2_* values found across 105 vertebrate populations in a meta-analysis of 50 HFC studies (average of 0.007) [22], and on the same order as ~60% of the local populations surveyed in a long-lived tree [23]. Our estimate of significant variation in heterozygosity provides the opportunity to examine potential drivers of this variance and examine how density, spatial structure and dispersal contribute to a non-uniform distribution of heterozygosity.

In our study, beyond simply estimating identity disequilibrium, we use two types of simulation to explore how demography shapes variation in inbreeding. The first simulation shows how the spatial pattern of pollen dispersal affects the distribution of heterozygosity. Simulated matings with the empirically measured leptokurtic pollen dispersal curve were consistent with the actual *g_2_*, compared to matings with nearest neighbors or a Gaussian pollen dispersal. This result is somewhat surprising because we did not include the complexities of the mating system of *A. majus*. *Antirrhinum majus* has a gametophytic self-incompatibility system (GSI [24]), whereby the pollen detoxifies secretions from the style unless the pollen and style genotypes share alleles at the S-locus [25]. This system not only prevents selfing, but also reduces mating among relatives (i.e., biparental inbreeding) because related plants are more likely to share S-alleles [4], [26]. Thus, we might expect that our simulated matings would have lower mean heterozygosity than the empirical measurement; yet we found no evidence for such an effect. Indeed, we found that the excess of individuals with low heterozygosity, around half the mean, can be explained largely by a small amount of bi-parental inbreeding with leptokurtic pollen dispersal (Fig. 3, SM1.3: Fig. S2). However, we have little statistical power to distinguish this from rare selfing, which can occur in self-incompatible species.

Our second simulation approach asked whether heterogeneous density promotes variation in inbreeding, given strong fine-scale population structure indicated by a rapid decay in pairwise F_ST_ (over a few metres, Fig. 2A). Indeed, simulated pedigrees with uniformly distributed plants gave less identity disequilibrium than we observed. In contrast, simulated pedigrees conditioned on the actual, heterogeneous density of plants were consistent with identity disequilibrium measured in the field. This indicates that patchiness, combined with leptokurtic dispersal shapes the distribution of heterozygosity. Simulations with heterogeneous density also better capture empirical isolation-by-distance patterns than those with a uniform density (Fig 4B, SM1.4: Fig. S8). However, the effective population size of 15,500 individuals in the heterogeneous-density simulations is an order of magnitude larger than the average number of plants observed in a year (~2500). We believe that most plants are sampled each year, so that this discrepancy is more likely to be due to a seed bank, which is expected to substantially increase the effective population size [27]. Nevertheless, despite simplifications such as non-overlapping generations, no seed bank, and a simple SI system, the heterogeneous-density simulation accurately captures patterns of identity disequilibrium and isolation-by-distance.

Our estimation of identity disequilibrium illustrates a general problem with statistical comparisons in evolutionary biology. There are three sources of error in estimating *g*_2_: firstly, error generated from sampling a limited number of individuals, secondly, from sampling a limited number of SNPs, and thirdly from random variation between evolutionary realizations or trajectories. In our study, the first source (a limited number of individuals) is shown by the confidence intervals in Fig. 5, obtained by bootstrapping across individuals [18]. The second source of error (a limited number of SNPs) is shown by the substantial variation in *g*_2_ of the ten replicates of each of five pedigrees. Here, variation is generated by random meiosis amongst unlinked markers on a fixed pedigree. This variation could be reduced by increasing the number of SNPs, but the effective number of segregating sites that can be included in the analysis is fundamentally limited by the length of the genetic map. Finally, there is additional variation between pedigrees, due to the random assignment of parents in the simulations, which generates a random pedigree. The wide variation in estimates of *g*_2_ due to random meiosis, and to the random generation of the pedigree (Fig. 5) is an important reminder that estimates of parameters are typically limited by the randomness of evolution. The stochasticity of evolution can potentially generate error variance far higher than that due to the limited number of individuals or SNPs sampled.

In addition to analyzing the effect of population structure on the distribution of heterozygosity, our study highlights the potential of utilizing multiple statistics to estimate population structure. We have shown that the variance of heterozygosity due to identity disequilibrium can distinguish alternative dispersal and density distributions, which implies that in combination with pairwise F_ST_ as a function of distance, *g*_2_ can help estimate the demography. Genetic data contain far more information than is described by F_ST_ and *g*_2_; for example, the mean squared disequilibrium can be used to estimate effective population size [28], [29], and this extends naturally to the covariance of pairwise linkage disequilibrium as a function of distance. We could simply use a set of such statistics to inform demographic inference via ABC [30]. However, our preference would be to first develop a theoretical understanding of how realistic demographies influence statistical measures of spatial covariance in allele frequency, identity disequilibria, and linkage disequilibria.

The distribution of heterozygosity has often been measured to estimate inbreeding depression and examine correlation with fitness. Yet, this type of data has rarely been used to investigate population structure *per se* and as a complement to the more widely used pairwise identity, F_ST_. By bringing together local inbreeding and isolation-by-distance, our study provides a novel assessment of how dispersal and population density can explain both pairwise identity and the distribution of heterozygosity in spatially continuous populations. However, we have only begun to investigate how the distribution of heterozygosity can be shaped by population structure and demographic parameters. Our future work will focus on understanding how other features such as a seed bank influence genetic diversity, with the ultimate goal of deriving information about demographic history from the distribution of heterozygosity in populations that have fewer measured parameters. New models that include these complexities, as well as ecological, mating system and life history factors are required to extend our understanding of the drivers of population structure in natural populations.

## Acknowledgments

We thank the many volunteers and friends who have contributed to data collection in the hybrid zone over the years, in particular those who have managed field seasons: Barbora Trubenova, Maria Clara Melo, Tom Ellis, Eva Cereghetti, Lenka Matejovicova, Beatriz Pablo Carmona. Frederic Ferrer and Eva Salmerón Mateu have been immensely helpful with logistics at our informal field station, El Serrat de Planoles. We thank Sean Stankowski for technical help in producing figure 1. Part of this work was funded by Marie Curie COFUND Doctoral Fellowship and Austrian Science Fund FWF (grant P32166). This research was also supported by the Scientific Service Units (SSU) of IST Austria through resources provided by Scientific Computing (SciComp).

