## Supplemental Material 1 for "Effects of fine-scale population structure on the distribution of heterozygosity in a long-term study of *Antirrhinum majus*"

### Heading order follows the Methods and Results of the main text.

#### SM1.1 SNP panel

##### *Detailed Methods: SNP Panel*

For each individual, DNA was extracted from leaf material collected from the field site, and was genotyped for the SNP panel by LGC Genomics (Middlesex, UK) using the KASP genotyping platform. Due to repeated sampling of the same individuals across years, the error rate of this method could be calculated, and was found to be low (mean error rate < 0.1% per locus).

Candidate loci were identified using a draft *A. majus* reference genome (~ 630 Mb across eight linkage groups; courtesy of Yongbiao Xue, Beijing Institute of Genomics); see ref [13]. In this study, SNPs were chosen to have overall mean frequency between 0.1 and 0.9; 90% had frequency between 0.25 and 0.75. SNPs that showed excessive geographic differentiation were eliminated by requiring a linear regression gradient of allele frequency on east-west distance to be less than 0.09 $km^{-1}$; 90% of chosen SNPs had a gradient < 0.03 $km^{-1}$. Furthermore, we required that $F_{ST}<0.1$; $F_{ST}$ was calculated by dividing the region into 200m squares, yielding 164 non-empty demes. Finally, the overall heterozygote deficit, $F_{IS}$, was required to be between -0.1 and 0.2; 90% of chosen SNPs had $-0.04 < F_{IS} < 0.1$. SNPs with heterozygote deficit $F_{IS} > 0.2$ also showed high $F_{ST}$ and/or clinal gradient, whilst those with $F_{IS} < -0.1$ were likely due to genotyping artefacts (e.g., primers binding to more than one site in the genome). After applying these filters, 170 SNPs remained. Finally, we chose to work with the 91 SNPs that were assayed for at least 60% of the Planoles sample (i.e., at least 13,411 individuals).

#### SM1.2 Variation in inbreeding

##### *Detailed Methods and Results*

The identity disequilibrium that we find is due partly to associations between linked SNP, and partly to associations between unlinked SNP (73% vs. 27%, respectively). Table S1 shows that correlations in H between SNP within linkage groups are consistently positive, averaging Pearson's $r = 0.01126$ - much higher than the average correlation of 0.00240 between all pairs of SNP, which are mostly unlinked. For the 155 individuals with H<0.3, the mean correlation within linkage groups, 0.0354, is much higher, reflecting the shared inheritance of large blocks of genome for close relatives. Correlations are higher between adjacent SNP, and yet higher in highly inbred individuals.

**Table S1.** Correlations in heterozygosity within the 8 linkage groups (LG). The third and fourth columns give the mean correlation in H between loci within each linkage group, for all 22,353 individuals versus for the 155 individuals with H<0.3. The last two columns give the mean correlations between adjacent SNPs. Note that 1 of the 91 SNP was not assigned to a linkage group.

| | | within LG | | adjacent SNP | |
|---|---|---|---|---|---|
| LG | No. SNPs | all inds | H<0.3 | all inds | H<0.3 |
| 1 | 13 | 0.01828 | 0.04423 | 0.08998 | 0.15209 |
| 2 | 15 | 0.02929 | 0.05021 | 0.12139 | 0.13723 |
| 3 | 10 | 0.00459 | 0.03298 | 0.00958 | 0.06359 |
| 4 | 12 | 0.00292 | -0.00370 | 0.00294 | -0.00530 |
| 5 | 12 | 0.02407 | 0.07854 | 0.02654 | 0.10365 |
| 6 | 15 | 0.00402 | 0.03128 | 0.00982 | 0.06743 |
| 7 | 6 | 0.00388 | 0.03862 | -0.00159 | 0.03416 |
| 8 | 7 | 0.00304 | 0.01089 | 0.00334 | 0.00805 |
| Mean | 90 | 0.01126 | 0.03538 | 0.03275 | 0.07011 |

#### SM1.3 Effects of pollen dispersal on heterozygosity

##### *Detailed Methods and Results*

The distribution of heterozygosity of offspring depends on distance between parents. We show this by simulating offspring, using all field-sampled individuals as mothers (Mathematica notebook in electronic supplementary material [16]). We chose fathers close to a given distance away, by choosing 12 points evenly spaced on a circle, and taking the nearest individual to any of those points. The genotype of the offspring was determined by Mendelian inheritance based on parental genotypes. The mean heterozygosity of offspring from two parents is linearly related to their pairwise identity; thus, the increase in mean identity with distance (Fig. S1A,C) is a precise reflection of the decay in $F_{ST}$. The variance in heterozygosity decreases with distance, as individuals become less related (Fig. S1B,D). Both mean and variance of H change sharply over scales of a few metres, and are hardly affected by linkage (compare gray and black lines in Fig. S1). The observed values (horizontal lines in Fig. S1) are consistent with pollination from fathers ~10m away, but are of course the product of a broad distribution of distances.

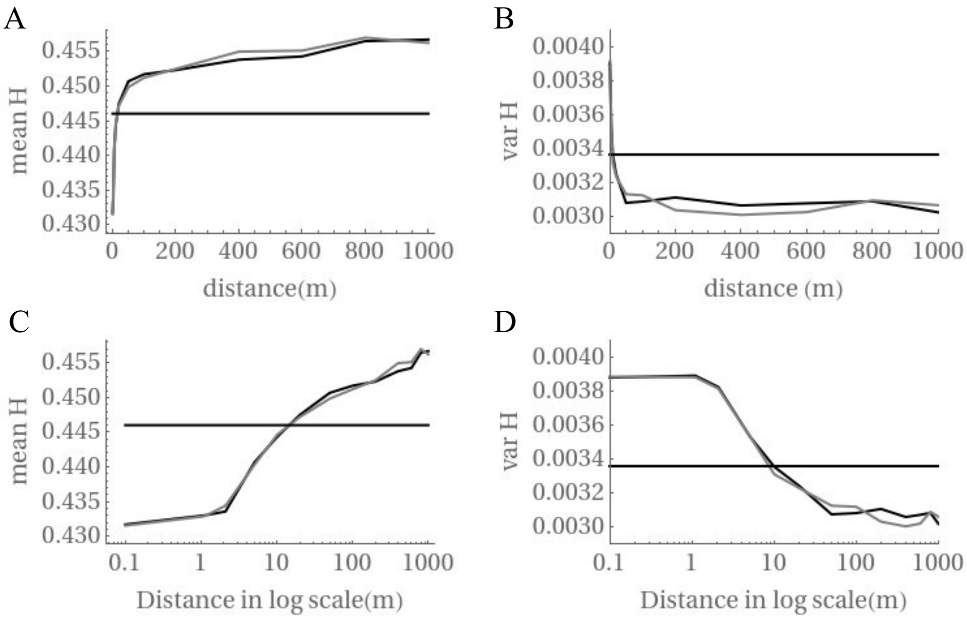


**Figure S1.** Mean (A, C) and variance (B,D) of H as a function of the distance between parents. Offspring are generated with no linkage (black) or with linkage (gray); observed values are shown as a horizontal line. Plots C and D show distance in a log scale.

**Table S2.** Mean and variance of multilocus heterozygosity (H) and identity disequilibrium ($g_2$) from field data and offspring simulated from three possible patterns of pollen dispersal (a leptokurtic kernel, a Gaussian kernel, and pollen from the nearest neighbour).

| | H mean | H variance | $g_2$ | $g_2$ CI |
|---|---|---|---|---|
| Field data | 0.4460 | 0.0034 | 0.0029 | 0.0026 - 0.0033 |
| Leptokurtic offspring | 0.4458 | 0.0034 | 0.0020 | 0.0016 - 0.0024 |
| Gaussian offspring | 0.4323 | 0.0039 | 0.0053 | 0.0049 - 0.0057 |
| Neighbour offspring | 0.4314 | 0.0039 | 0.0056 | 0.0051 - 0.0060 |

**Table S3.** Test statistic and *p*-value from *t*-test, *F*-test and Kolmogorov-Smirnov (KS) test for each pairwise comparison between heterozygosity calculated from field data and offspring simulated from leptokurtic, Gaussian, and nearest neighbour matings.

| | | *t*-test | | *F* test | | KS test | |
|---|---|---|---|---|---|---|---|
| Dataset | Dataset | *t* | *p* | *F* | *p* | *D* | *p* |
| Field | Leptokurtic | 1.08 | 0.281 | -0.55 | 0.579 | 0.015 | 0.015 |
| Field | Gaussian | 24.06 | <0.001 | -10.02 | <0.001 | 0.094 | <0.00001 |
| Field | Neighbour | 25.17 | <0.001 | -9.78 | <0.001 | 0.103 | <0.00001 |
| Leptokurtic | Gaussian | 23.07 | <0.001 | -9.80 | <0.001 | 0.086 | <0.00001 |
| Leptokurtic | Neighbour | 24.19 | <0.001 | -9.36 | <0.001 | 0.090 | <0.00001 |
| Gaussian | Neighbour | 1.06 | 0.29 | 0.31 | 0.75 | 0.009 | 0.36 |

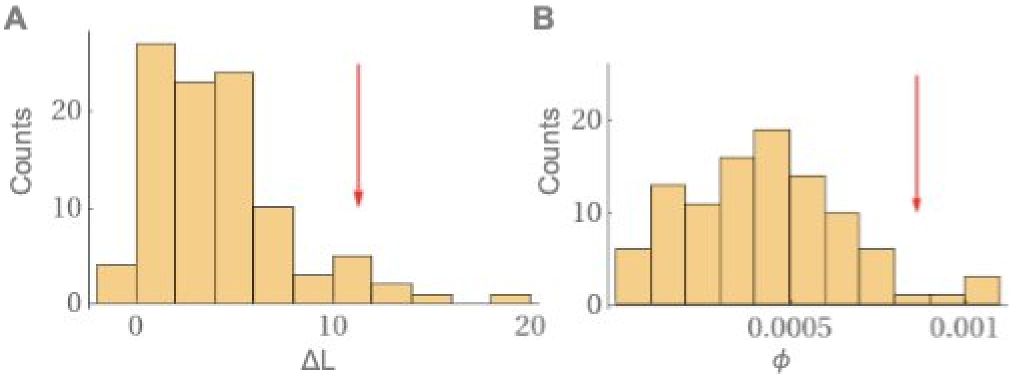


**Figure S2:** The distribution of increase in the likelihood (ΔL) (A) and selfing rate, ϕ (B) between the single and mixed Gaussian distributions from 100 replicates of simulated matings from leptokurtic dispersal distribution. The red arrow points to the value observed in the field data.

#### SM1.4 Heterozygosity in a simulated spatial pedigree

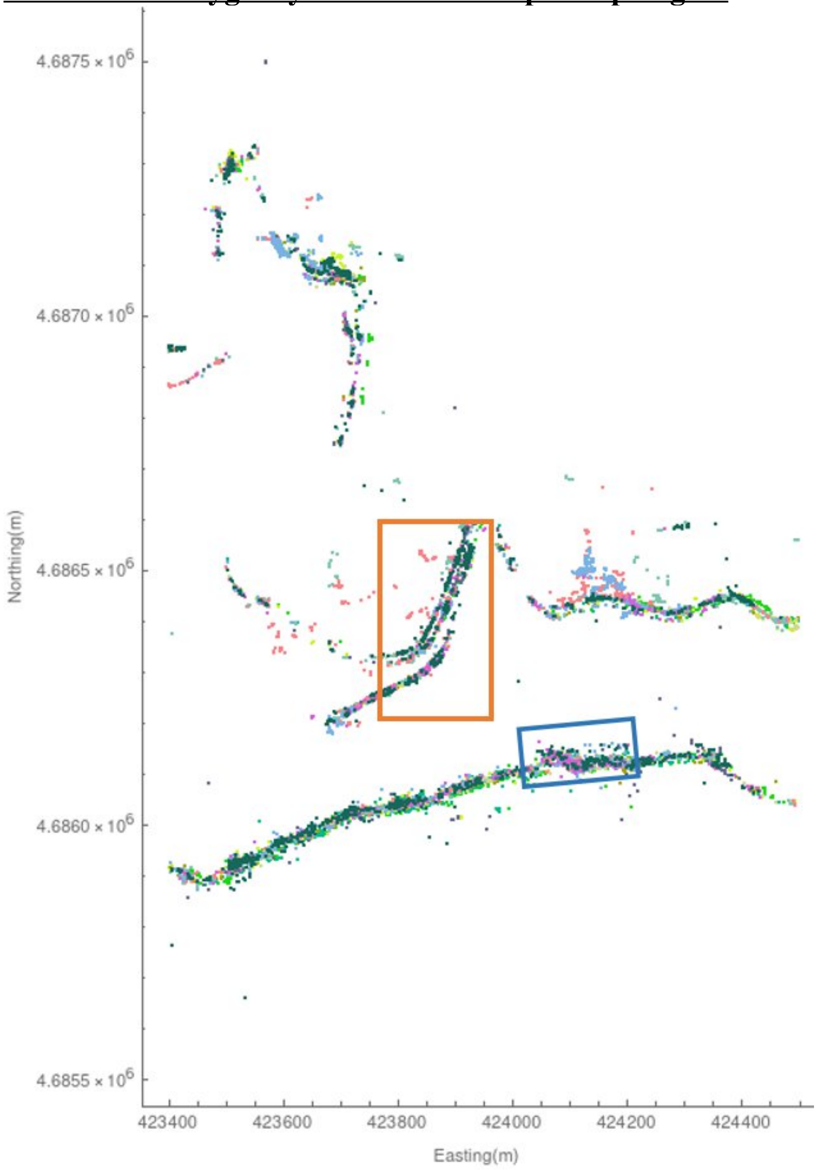


**Figure S3.** Individual plant locations in the simulated region of the field site. Each colour represents a different year. See Fig. S2 for time series of boxed areas.

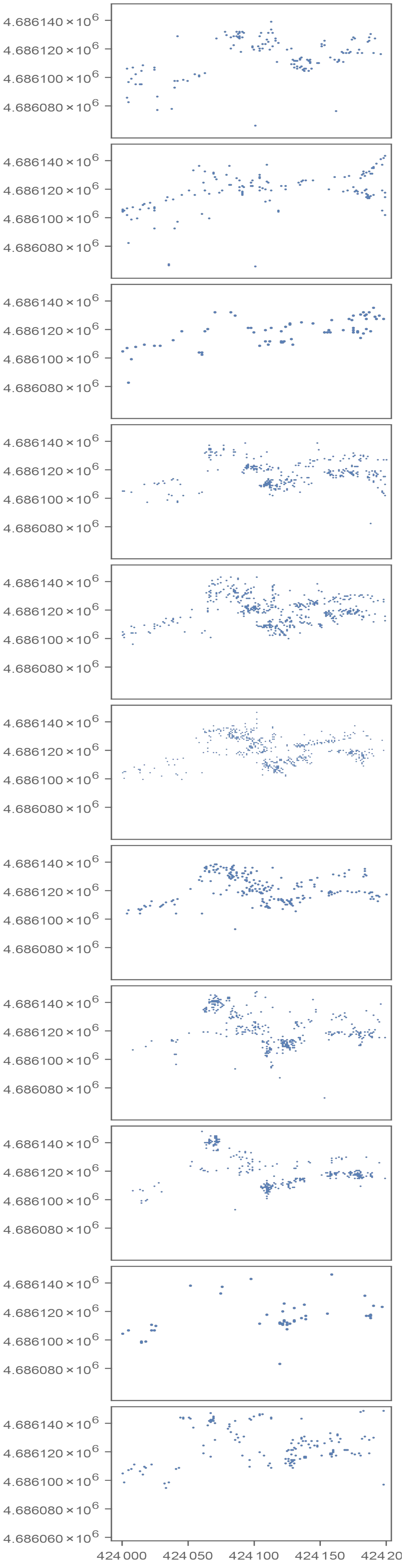
4.686140 × 10^6
4.686120 × 10^6
4.686100 × 10^6
4.686080 × 10^6
4.686060 × 10^6
424 000
424 050
424 100
424 150
424 20

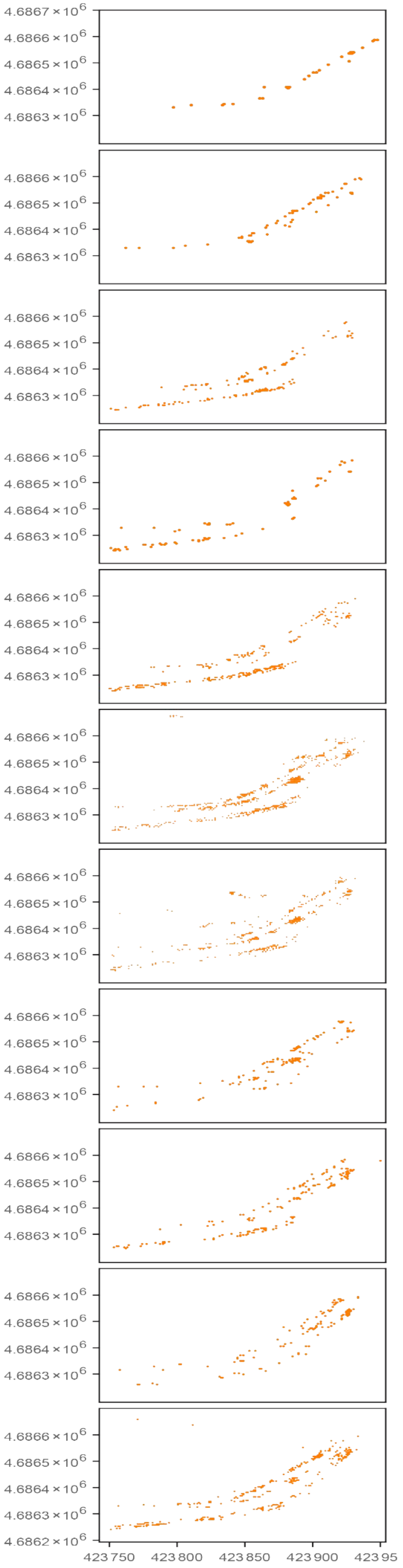
4.6867 × 10^6
4.6866 × 10^6
4.6865 × 10^6
4.6864 × 10^6
4.6863 × 10^6
4.6862 × 10^6
423 750
423 800
423 850
423 900
423 95

101

**Figure S4.** Close-ups of sections of the lower road (left) and upper road (right) from the field data, showing changes in patchiness over time from 2009 (top) to 2019 (bottom). Sections are denoted as blue and orange in Fig. S3.

##### *Detailed Methods and Validations for Simulated Spatial Pedigree*

The simulation was validated by comparing with the analytical results for the rate of increase of average identity by descent with time in a panmictic population (simulated by choosing parents without regard to their locations). It was also validated for a stepping stone model (simulated by locating individuals in discrete demes on a toroidal grid, and allowing dispersal only between nearest neighbors) by comparing rate of change of average identity by descent with average identity by state. The agreement was close for both comparisons (Fig S5; colors cannot be distinguished, as the curves overlap).

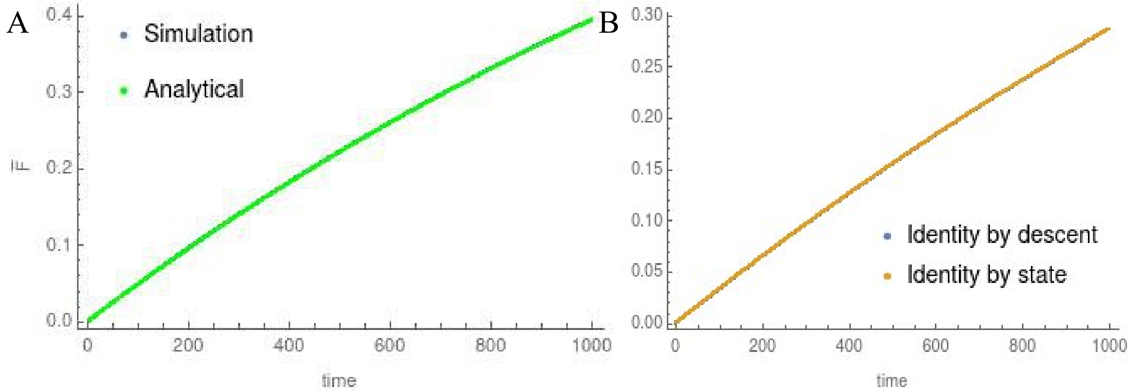


**Figure S5**. Rate of increase of average pairwise identity ($\bar{F}$) with time for a panmictic population (A) and stepping stone model on a torus (B).

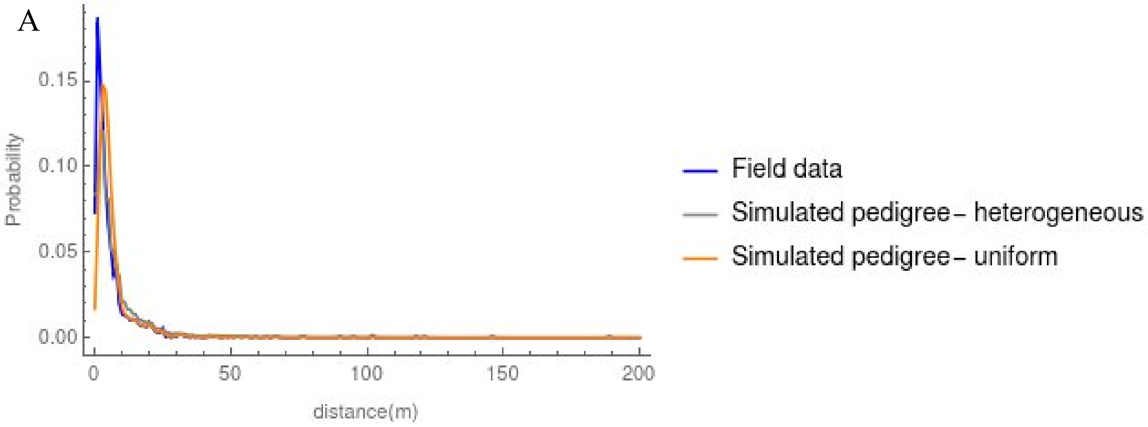

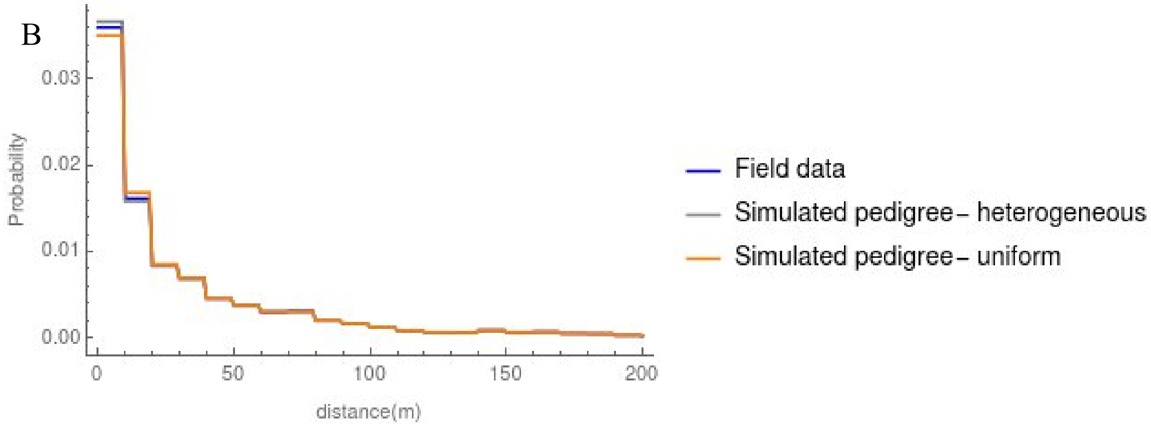


**Figure S6.** Realized (gray and orange) and proposed (blue) seed (A) and pollen (B) dispersal distribution for the simulated pedigrees with heterogeneous and uniform population structure. Due to computational constraints, these are calculated from the last 300 generations for the simulated pedigree with uniform density. The pedigree with $F_{ST}$ closest to that of the field data is shown for the heterogeneous case (also in Fig S7B, S8).

**Table S4.** Mean and standard deviation (SD) of the proposed and realized seed and pollen dispersal distributions for the simulated pedigrees with uniform and heterogeneous spatial structure.

| | Seed dispersal | | Pollen dispersal | |
|---|---|---|---|---|
| | Mean | SD | Mean | SD |
| Proposed | 9.52468 | 38.2242 | 62.684 | 119.327 |
| Heterogeneous pedigree 1 | 12.797 | 40.1884 | 62.5955 | 118.765 |
| Heterogeneous pedigree 2 | 12.7785 | 40.0344 | 62.6493 | 118.834 |
| Heterogeneous pedigree 3 | 12.7724 | 40.0098 | 62.5716 | 118.682 |
| Heterogeneous pedigree 4 | 12.7863 | 40.0749 | 62.6127 | 118.778 |
| Heterogeneous pedigree 5 | 12.7858 | 40.0773 | 62.5902 | 118.774 |
| Heterogeneous- average | 12.784 | 40.0770 | 62.6039 | 118.767 |
| Uniform pedigree | 10.4706 | 38.0741 | 63.1388 | 119.122 |

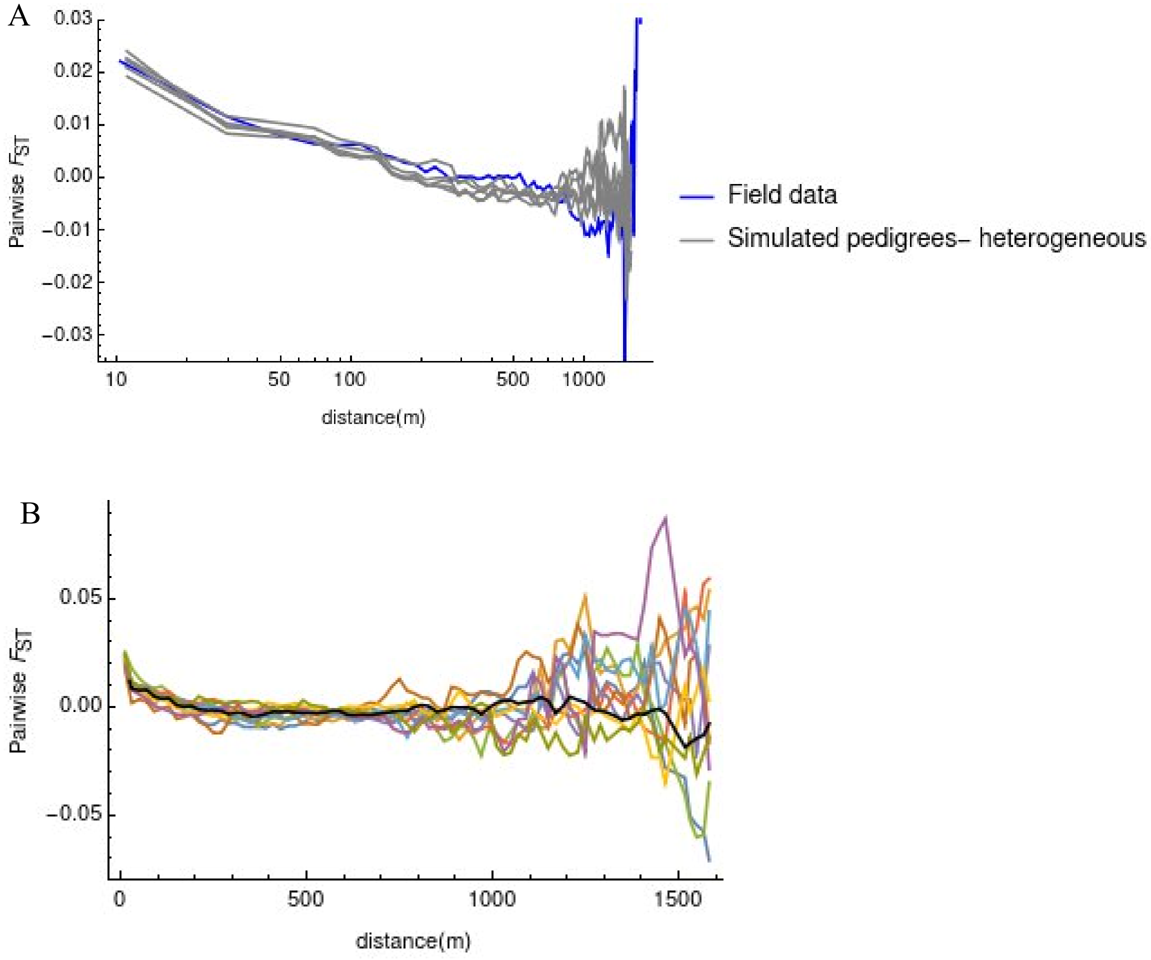


**Figure S7.** (A) Isolation by distance for the field data (blue) and five simulated population pedigrees (gray) plotted on a log scale. (B) Isolation by distance from ten replicates of a single pedigree along with their average shown in black.

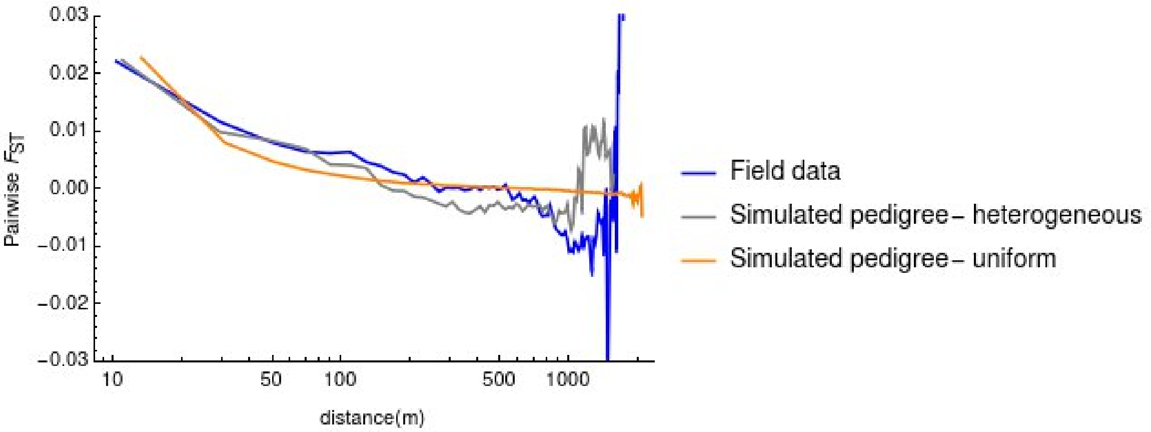


**Figure S8.** Isolation by distance for the field data (blue), simulated pedigree with realistic spatial structure (gray) and uniform density (orange) plotted on a log scale.

**Table S5**. $F_{ST}$, $F_{IS}$ and $g_2$ values from the field data and from simulated (sim.) pedigrees with heterogeneous and uniform density. For the simulations, mean ± standard deviation of ten replicate sets of genotypes are shown for each pedigree, across the five pedigree means, and across all 50 replicates (ten replicates for five pedigrees).

| | | $F_{ST}$ | $F_{IS}$ | $g_2$ |
|---|---|---|---|---|
| Field data | | 0.022 | 0.0211 | 0.00262 |
| Sim. heterogeneous | Pedigree 1 | 0.0192 ± 0.00383 | 0.0216 ± 0.00236 | 0.00274 ± 0.000723 |
| | Pedigree 2 | 0.0226 ± 0.00348 | 0.0254 ± 0.00179 | 0.00258 ± 0.000560 |
| | Pedigree 3 | 0.0222 ± 0.00221 | 0.0247 ± 0.00149 | 0.00312 ± 0.001120 |
| | Pedigree 4 | 0.0239 ± 0.00143 | 0.0244 ± 0.00132 | 0.00240 ± 0.000854 |
| | Pedigree 5 | 0.0208 ± 0.00346 | 0.0259 ± 0.00160 | 0.00235 ± 0.000448 |
| | Across pedigree means | 0.0217 ± 0.0029 | 0.0244 ± 0.0017 | 0.00264 ± 0.000741 |
| | Across all 50 replicates | 0.0217 ± 0.0033 | 0.0244 ± 0.0023 | 0.00264 ± 0.000797 |
| Sim. uniform | Pedigree 1 | 0.0226 ± 0.0009 | 0.0203 ± 0.00059 | 0.00171 ± 0.000083 |
